# A glucosinolate-rich beverage lowers blood lactate concentrations during submaximal exercise

**DOI:** 10.1101/2025.04.15.648889

**Authors:** Fredrik Vinge, Emma Tillqvist, Oscar Horwath, William Apró, Filip J Larsen, Michaela L Sundqvist

## Abstract

Glucosinolate-rich broccoli sprouts combined with intense exercise training for 7 days have been shown to reduce blood lactate concentrations during exercise, attenuate hypoglycemic events, improve physical performance, and reduce markers of oxidative stress. This study aimed to investigate the acute, dose-dependent effects of glucosinolate-rich red kale sprouts (GRS) on blood lactate and blood glucose following the ingestion of three different doses.

Fifteen healthy participants consumed 37.5 g or 75 g of GRS or an isocaloric placebo blended into a beverage on three separate occasions. The participants cycled on an ergometer at three submaximal work rates before and three hours after ingestion. Measurements of oxygen uptake, substrate-level oxidation, blood lactate, blood glucose, and ratings of perceived exertion were taken before and after each cycling interval.

The intake of glucosinolate-rich sprouts acutely decreased blood lactate levels during submaximal cycling and increased blood glucose levels at rest. The largest reduction in blood lactate was observed at the 37.5 g dose compared to placebo, where the concentration was 0.4 ± 0.2 mM lower at work rate close to threshold (p = 0.003). For the 75 g dose, the reduction in blood lactate was 0.25 ± 0.1 mM (p = 0.02). No significant effects were seen in the lowest work rate. The mean resting glucose level was 3.9 ± 0.1 mM following placebo, compared to 4.3 ± 0.1 mM after intake of either 37.5 g or 75 g dose (p < 0.01, respectively).

These findings suggest that glucosinolate-rich broccoli sprouts have a lactate-lowering effect during submaximal efforts, which may have important implications for supplementation for improving endurance performance.

## BACKGROUND

In endurance exercise, lactate accumulation is closely correlated with endurance performance (Sjödin & Svedenhag, 1985). The lactate threshold represents the intensity at which lactate begins to accumulate rapidly in the blood. Typically, the lactate threshold occurs around 4 mM (Billat, 1996), though there is considerable individual variation. Lower lactate levels at submaximal exercise intensities are a hallmark of positive adaptations to a period of aerobic endurance exercise training (Hurley et al., 1984). Recently, regular lactate measurements have gained a central role in advancing training models, such as lactate-guided threshold interval training (LGTIT). This approach tailors training intensity based on an athlete’s blood lactate levels and has shown beneficial results when implemented in elite runners (Casado et al., 2023).

To lower lactate levels during exercise, different pharmacological approaches have also been explored. One very recognized attempt was the use of dichloroacetate (DCA), a drug that stimulates pyruvate dehydrogenase activity. Activation of pyruvate dehydrogenase by DCA promotes the utilization of pyruvate in the mitochondria, thereby decreasing lactate accumulation. While DCA consistently reduces lactate levels during submaximal cycling, its effectiveness in improving exercise performance seems to be context-dependent (Calvert et al., 2008; Carraro et al., 1989; Timmons et al., 1998).

Recently, a specific group of phytochemicals, isothiocyanates, have gained attention for their ability to alter metabolic responses and gene expression during exercise. Isothiocyanates, naturally found in cruciferous vegetables like kale, broccoli, and cabbage, are derived from the hydrolysis of glucosinolates, a reaction catalyzed by the enzyme myrosinase (Fahey et al., 1997). Supplementation with the most well-studied isothiocyanate, sulforaphane (SFN), has been shown to activate the transcription factor Nuclear Factor Erythroid 2-Related Factor 2 (Nrf2) (Oh et al., 2017). Nrf2 is recognized as the primary regulator of cellular oxidative stress and has been identified as a key coordinator of the adaptive response to exercise training (Vargas-Mendoza et al., 2019). In a study by Oh et al. (2017), mice receiving intraperitoneal injections of SFN for 3 days prior to exhaustive exercise exhibited reduced oxidative stress markers and improved running capacity. Also, treatment with SFN in animal exercise studies under hypoxic conditions resulted in lower plasma lactate levels, increased expression of monocarboxylate transporters (MCTs), and enhanced lactate dehydrogenase (LDH) activity which facilitate the conversion of lactate to pyruvate (Wang et al., 2020). Furthermore, SFN treatment during a 6-week high-intensity interval training (HIIT) protocol in mice improved the antioxidative capacity of skeletal muscle, in response to acute exhaustive exercise, by increasing protein expression of Nrf2 (Wang et al., 2022).

Our research group recently conducted a human trial in which participants consumed 75 g of glucosinolate-rich broccoli sprouts (GRS) twice daily for 7 consecutive days, during a period of high-intensity interval training (HIIT). The results showed a reduction in oxidative stress markers, decreased lactate levels during submaximal cycling, and enhanced physical performance compared to the placebo treatment (Flockhart et al., 2023).

This single-blinded, randomized, placebo-controlled crossover study aimed to assess the acute effects of GRS on lactate and glucose metabolism during submaximal cycling, three hours after consuming varying doses (0 g, 37.5 g, or 75 g) of GRS.

## METHOD

### Study population

Participants were recruited through advertisements at the Swedish School of Sports and Health Sciences (GIH) and via a study-participant recruitment website (Accindi.com). Eligible participants were between 18-40 years old, without any chronic diseases. Exclusion criteria included diseases such as liver disease, heart disease, diabetes, and severe anxiety disorders. A total of 23 participants were screened, and 11 females and 4 men completed the study. Baseline characteristics are presented in Table 1.

**Table 1.** Baseline characteristics, mean ±SEM.

|  | **Female** | **Male** | **Total** |
| --- | --- | --- | --- |
| **n** | 11 | 4 | 15 |
| **Age (years)** | 27.6 ± 1.6 | 25.0 ± 1.1 | 26.9 ± 1.3 |
| **Body weight (kg)** | 63.9 ± 2.1 | 76.9 ± 1.9 | 67.4 ± 2.2 |
| **Height (cm)** | 167.3 ± 0.8 | 177.8 ± 1.6 | 170.1 ± 1.4 |
| **VO<sub>2</sub>max (ml/min<sup>-1</sup> kg<sup>-1</sup>)** | 38.5 ± 1.7 | 47.8 ± 3.3 | 40.9 ± 1.8 |

### Standardization of food intake and training

Participants were advised to maintain their normal diet but avoid consuming cruciferous vegetables (e.g., broccoli, kale) and nitrate-rich vegetables (e.g., lettuce, beetroot) three days before the laboratory visits. They were also instructed to keep a 24-hour dietary record the day before the first visit, and this recorded food intake was replicated for the subsequent test days to ensure standardized dietary intake. Participants were further instructed to refrain from caffeine intake on test days and to avoid any exercise training 36 hours before visits.

### Quantification of glucosinolates and isothiocyanates

Glucosinolate measurement was performed following the method of Mawlong et al., with minor modifications (Mawlong et al., 2017). Freeze-dried sprouts were homogenized in 80% methanol and then centrifuged at 3000 rpm for 4 minutes at room temperature, and the supernatant was collected. A 50 μL aliquot was used for analysis, mixed with 50 μL methanol and 250 μL of 2 mM STCP solution (prepared by dissolving 58.85 mg sodium tetrachloropalladate in 170 μL concentrated HCl and 100 mL dH_2_O). The mixture was thoroughly mixed and immediately analyzed via spectrophotometry at 425 nm. Sinigrin was used as the internal standard.

Isothiocyanate content (L-sulforaphane and iberin) was measured at the Sweden Metabolomics Center in Umeå with Liquid Chromatography-Mass Spectrometry (LC-MS) in four sprout varieties from cruciferous vegetables, including broccoli sprouts and red kale sprouts. Sample preparation was performed according to (A et al., 2005) with some modifications. Before LC-MS analysis, samples were re-suspended in methanol and water. Analysis was performed in both positive and negative ion modes using an Agilent 1290 Infinity UHPLC system (Agilent Technologies, Waldbronn, Germany) coupled with an Agilent 6546 Q-TOF mass spectrometer. Chromatographic separation was achieved on a C18 column with gradient elution. Reference ions were used for accurate mass calibration. Instrument parameters, including gas temperature, flow rates, and voltages, were optimized for detection. Data processing was conducted using Agilent MassHunter Profinder (Agilent Technologies Inc., Santa Clara, USA), with targeted feature extraction for isothiocyanate metabolites. Concentrations of isothiocyanates were determined based on calibration curves. A detailed description of the methods can be found in the supplemental material.

### Preparation of red kale sprout beverage

Red kale sprouts were selected for the intervention as they provided the highest levels of total glucosinolates and sulforaphane. Each bottle contained sprout concentrate equivalent to 37.5 g of fresh sprouts and in the GRS bottles’ this was equivalent to 606 mg glucosinolates and 8 mg of sulforaphane, 13 g carbohydrates, and ∼62 kcal. Accordingly, 75 g fresh sprouts contained 1212 mg glucosinolates, 16 mg sulforaphane 26 g carbohydrates, and ∼124 kcal. The PLA was virtually devoid of glucosinolates. To produce the study beverages the glucosinolate-rich, red kale sprouts (GRS) and the alfalfa sprouts (PLA) were pressed, strained, and mixed with water and black currant cordial. Subsequently, the beverage underwent low-temperature pasteurization and was bottled in 250 mL glass bottles. The GRS and PLA beverages used in the study were produced at Balsgård Foodtech, Sweden. The entire batch was frozen directly after production and thawed 30 minutes before use, ensuring consistent taste and glucosinolate level throughout the study period. To enhance the conversion of potentially remaining glucosinolates to isothiocyanates, 0.75 g of myrosinase-rich brown mustard seeds were mixed into the drink before consumption.

### Pre-tests

During the screening visit, participants were familiarized with cycling on the ergometer bike (PT1, Wattbike, Nottingham, UK), and their lactate threshold was determined after cycling at five different submaximal intensities for 5 min. A heart rate monitor (Polar Electro OY, Kempele, Finland) was attached to the chest during cycling and capillary blood was drawn from the fingertip, then analyzed with an automatic analyzer for glucose and lactate (Biosen C line, EKF-Diagnostics, Cardiff, UK). The submaximal cycling test was succeeded by an incremental VO_2_max test to determine maximal oxygen uptake. A facemask with a turbine was attached to participants during both submaximal- and maximal cycling to analyze oxygen consumption (Quark CPET, Cosmed Albano Laziale, Italy). Blood pressure (BP) measurements were taken in the left arm after 5 minutes of rest in a supine position. Three measurements, with 60 seconds between each, were performed using an automatic blood pressure monitor (M2, Omron, Kyoto, Japan). The average of the two last measurements was considered the subject’s BP.

### Dose-response test

On test days, participants arrived in the morning on three independent days, separated by 7 days. The exercise tests were conducted at the same time of day for all tests. Body weight and blood pressure measurements, along with resting capillary blood glucose and lactate, were taken. This was followed by cycling at three submaximal intensities. The resistance on each interval was based on the screening threshold test and aimed to reach 2 mM, 3 mM, and 4 mM lactate after 5 min cycling. The mean effect (Watt) for the three different submaximal workloads on the ergometer bike was 104 ± 44 W, 124 ± 46 W, and 140 ± 46 W. After completion of the three intervals, participants ingested a beverage containing either 0 g GRS (PLA), 37.5 g GRS (∼8 mg SFN), or 75 g GRS (∼16 mg SFN) together with 0.75 g of mustard seeds. Three hours after intake, participants repeated the cycling protocol followed by blood sampling. Myeloperoxidase was measured in EDTA-plasma using the myeloperoxidase Duo Set ELISA kit according to the manufacturer’s instructions (R&D Systems, Abingdon, UK). Between the pre-and post-test participants were fasting but allowed to drink water ad libitum.

### Statistical analysis

A linear mixed model was used to analyze the dose-dependent effects with the change in blood lactate and glucose from pre to post as response variables. The analysis was conducted using Python, with the Statsmodels library for statistical modeling. Comparison between doses was made using the placebo dose as the reference category. This categorization facilitated the comparison of treatment effects across different dosage levels. The model accounted for fixed effects of the dose, VO2max, and baseline levels of lactate and glucose, both at rest and under submaximal exercise. Additionally, each subject was treated as a random effect to account for the inter-subject variability. This approach allowed us to assess the effect of the treatment while controlling for individual differences and baseline metabolic states. A p-value below 0.05 was considered significant, and data are presented as mean ± SEM.

## RESULTS

### A glucosinolate-rich beverage lowers blood lactate during submaximal cycling

Consumption of GRS reduced the blood lactate concentration at a given absolute work rate compared to PLA. The largest reduction was observed at the 37.5 g dose indicating a U-shaped dose-response relationship. Compared to placebo the lactate concentration was 0.2 ± 0.1 mM lower at work rate 2 (p = 0.01) and 0.4 ± 0.2 mM lower at work rate 3 (p = 0.003). For the 75-gram dose, the reduction was 0.1 ± 0.1 mM at work rate 2 (p = 0.16) and 0.25 ± 0.1 mM at work rate 3 (p = 0.02). No significant effects were seen in the lowest work rate (Figure 1) or resting lactate in the morning compared to resting lactate after a 3-hour fasting rest.

**Figure 1.**
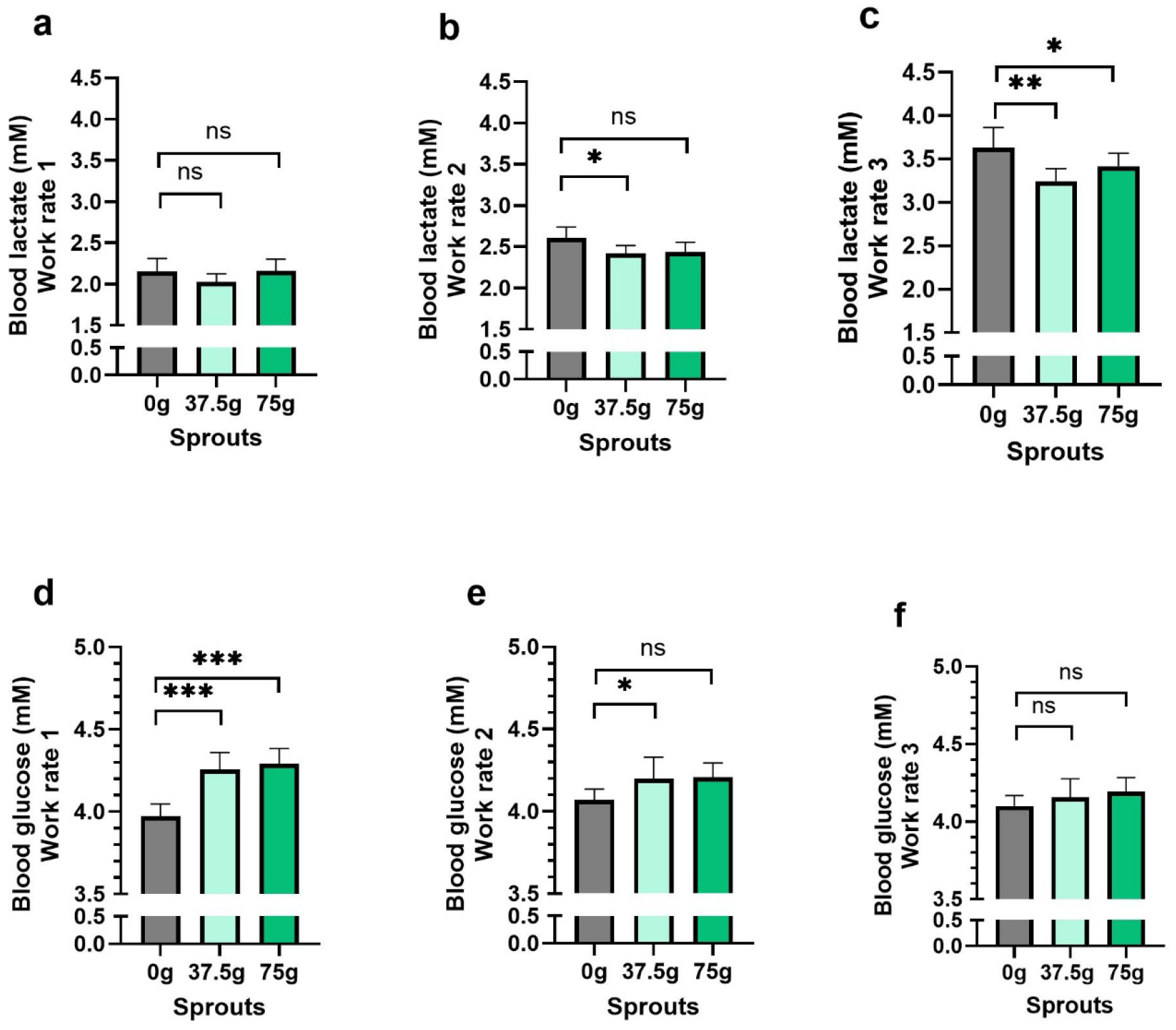
Capillary blood lactate levels (a-c) and capillary blood glucose levels (d-f) during cycling at three work rates on an ergometer bike, measured 3h after intake of a beverage containing 0 g, 37.5 g, or 75 g GRS. Data are presented as mean ± SEM, n = 15. * = p < 0.05, ** = p < 0.01, *** = p < 0.001.

### Blood glucose was normalized after ingesting a glucosinolate-rich beverage

During submaximal cycling, blood glucose after 37.5 g GRS was significantly lower both at work rate 1 (p = 0.01) and work rate 2 (p = 0.04) compared to PLA, but not at work rate 3 (p = 0.13). Resting blood glucose after ingesting 37.5 g and 75 g GRS were higher (p = <0.01, respectively) compared to PLA intake (Figure 2). Specifically, after 3 hours of rest, the mean glucose level was 3.9 ± 0.1 mmol/L following PLA, compared to 4.3 ± 0.1 mmol/L after intake of either 37.5 g or 75 g of GRS. In total 10 out of 15 participants were hypoglycemic (below 4 mM blood glucose) after PLA compared to 4 participants when supplementing with 37.5 g GRS and 2 participants when supplementing with 75 g. However, resting blood glucose was significantly higher in the morning compared to 3h rest in fasting (p < 0.0001 for PLA, p = 0.02 for 37,5g GRS and p = 0.004 for 75 g GRS).

**Figure 2.**
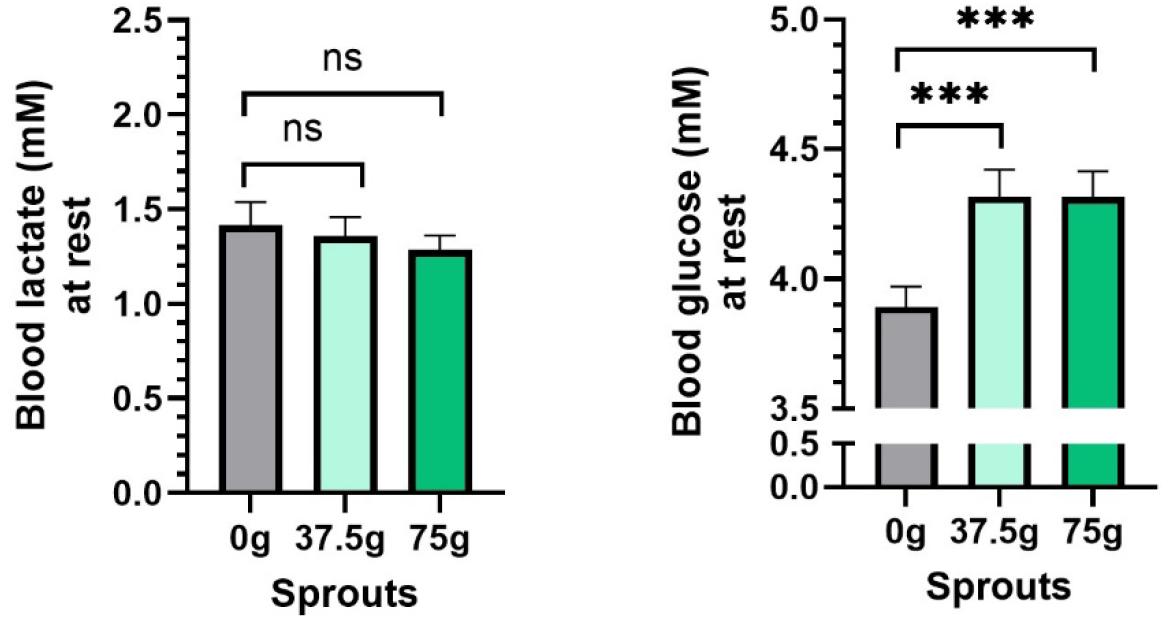
Capillary blood lactate (a) and blood glucose (b) levels during rest, measured 3h after intake of a beverage containing 0 g, 37.5 g, or 75 g of GRS. Data are presented as mean ± SEM, n = 15. *** = p < 0.001.

### A glucosinolate-rich beverage did not affect blood pressure or heart rate

There was no difference in resting systolic- or diastolic BP, or HR after intake of GRS compared to PLA (Figure 3).

**Figure 3.**
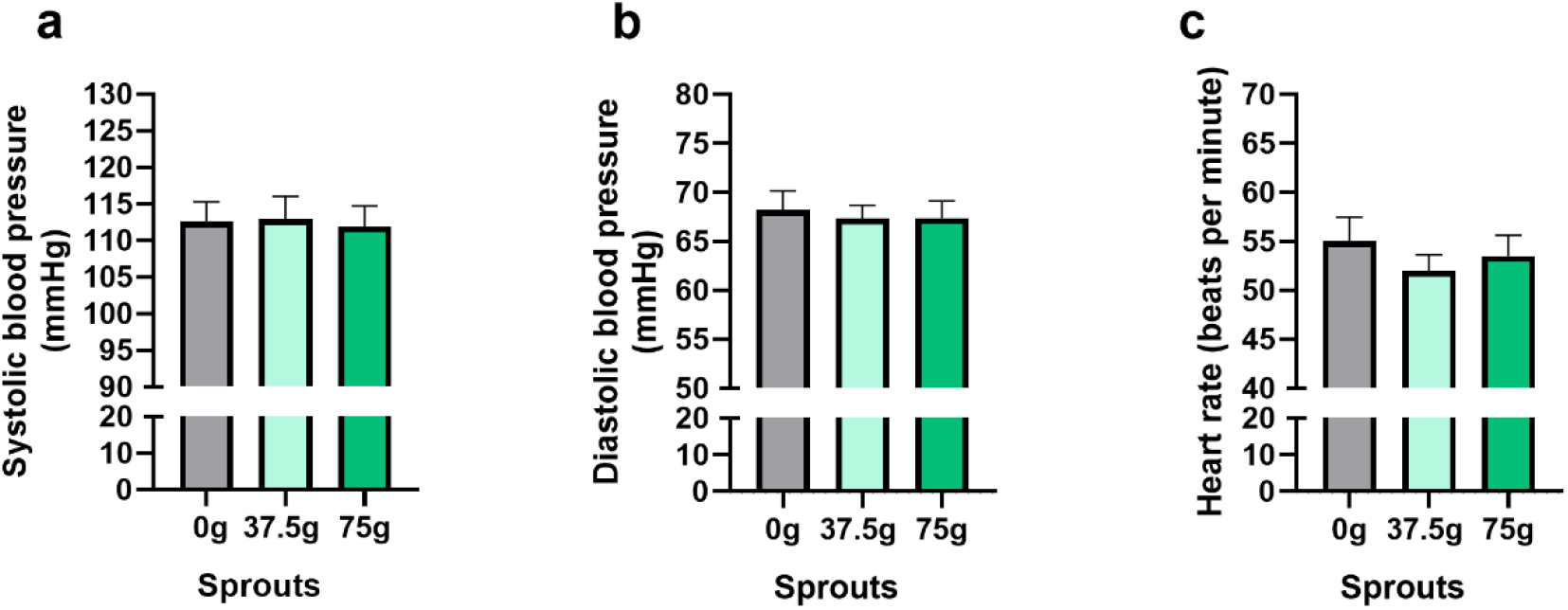
Resting systolic blood pressure (a), diastolic blood pressure (b), and heart rate (c), measured 3h after intake of a beverage containing 0 g, 37.5 g, or 75 g of GRS. Data are presented as mean ± SEM, n = 15. p < 0.05 is considered significant.

### Myeloperoxidase

To investigate any changes in oxidative stress markers after intake of GRS we measured myeloperoxidase concentrations in plasma. Mean myeloperoxidase levels were lower after 37.5 g and 75 g GRS (17.6 ± 6.1 and 17.8 ± 6.0 ng/mL, respectively), compared to PLA (19.3 ± 6.1 ng/mL) (Fig. 4).

**Figure 4.**
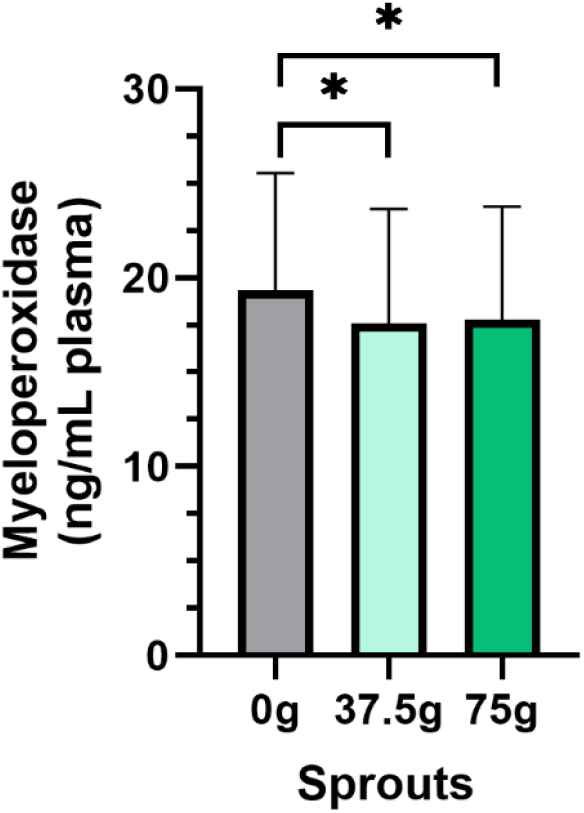
Mean myeloperoxidase levels 3h after intake of a beverage containing 0 g, 37.5 g, or 75 g GRS. Data are presented as mean ± SEM, n = 15. * = p < 0.05.

### Oxygen uptake, substrate use, and ratings of perceived exertion and heart rate

During work rate 1, the respiratory exchange ratio (RER) increased after the intake of 75 g of GRS, and after work rate 3, it was significantly reduced following the intake of 37.5 g of GRS. However, no differences in oxygen consumption or carbon dioxide production were observed (Figure 5). Additionally, respiratory frequency was slightly increased during work rate 3 after the intake of 37.5 g of GRS. No differences were observed in ventilation at any work rate or with the intake of PLA or GRS (Supplementary material, Figure. 1). Furthermore, there were no differences in the rating of perceived exertion between groups at any work rate (Supplementary material, Figure. 2).

**Figure 5.**
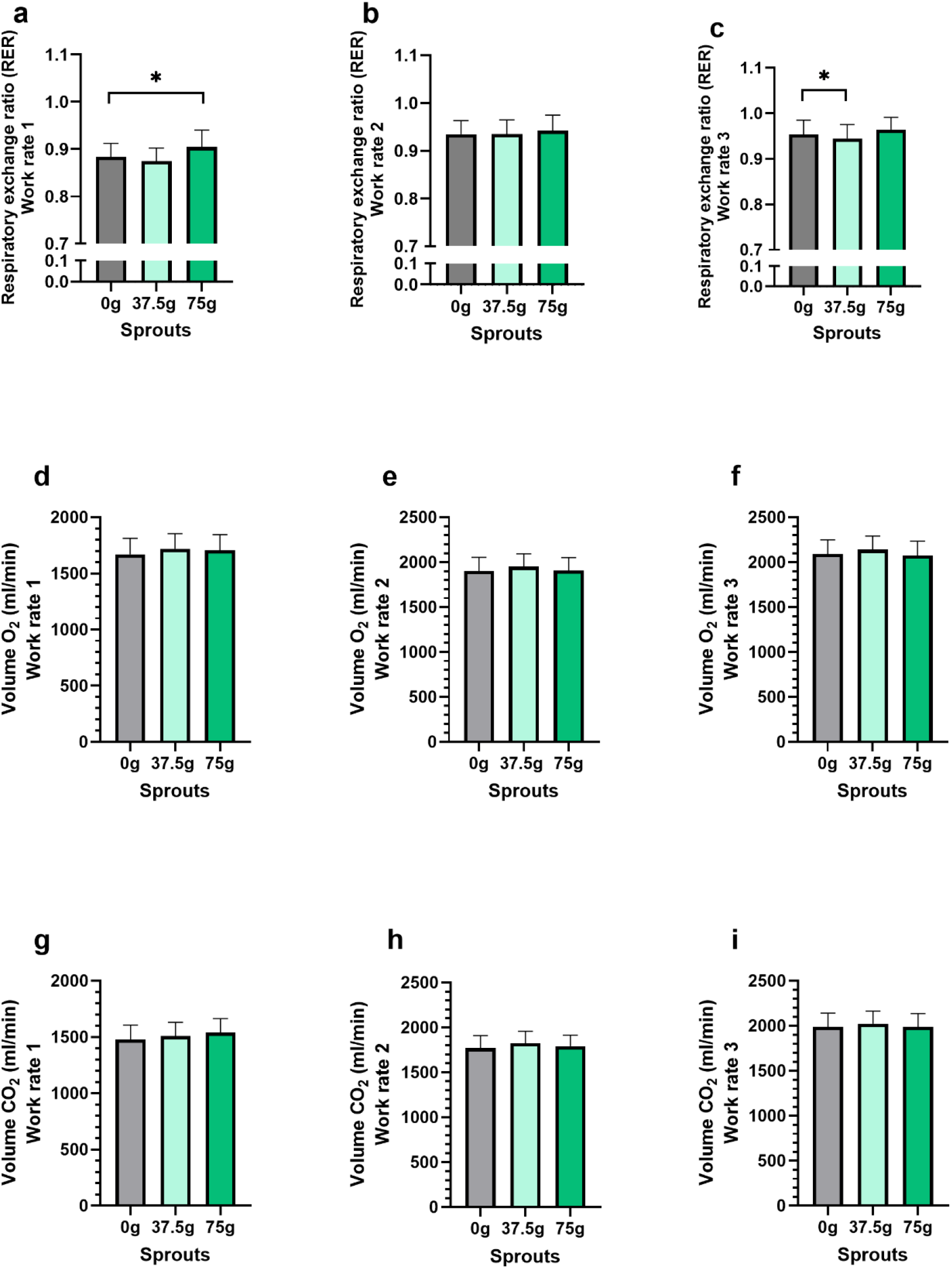
Respiratory exchange ratio (a-c), volume O_2_ consumed (d-f), and CO2 produced (g-i) during cycling at three work rates on an ergometer bike, measured 3h after intake of a beverage containing 0 g, 37.5 g, or 75 g GRS. Data are presented as mean ± SEM, n = 15. * = p < 0.05.

**Figure 6.**
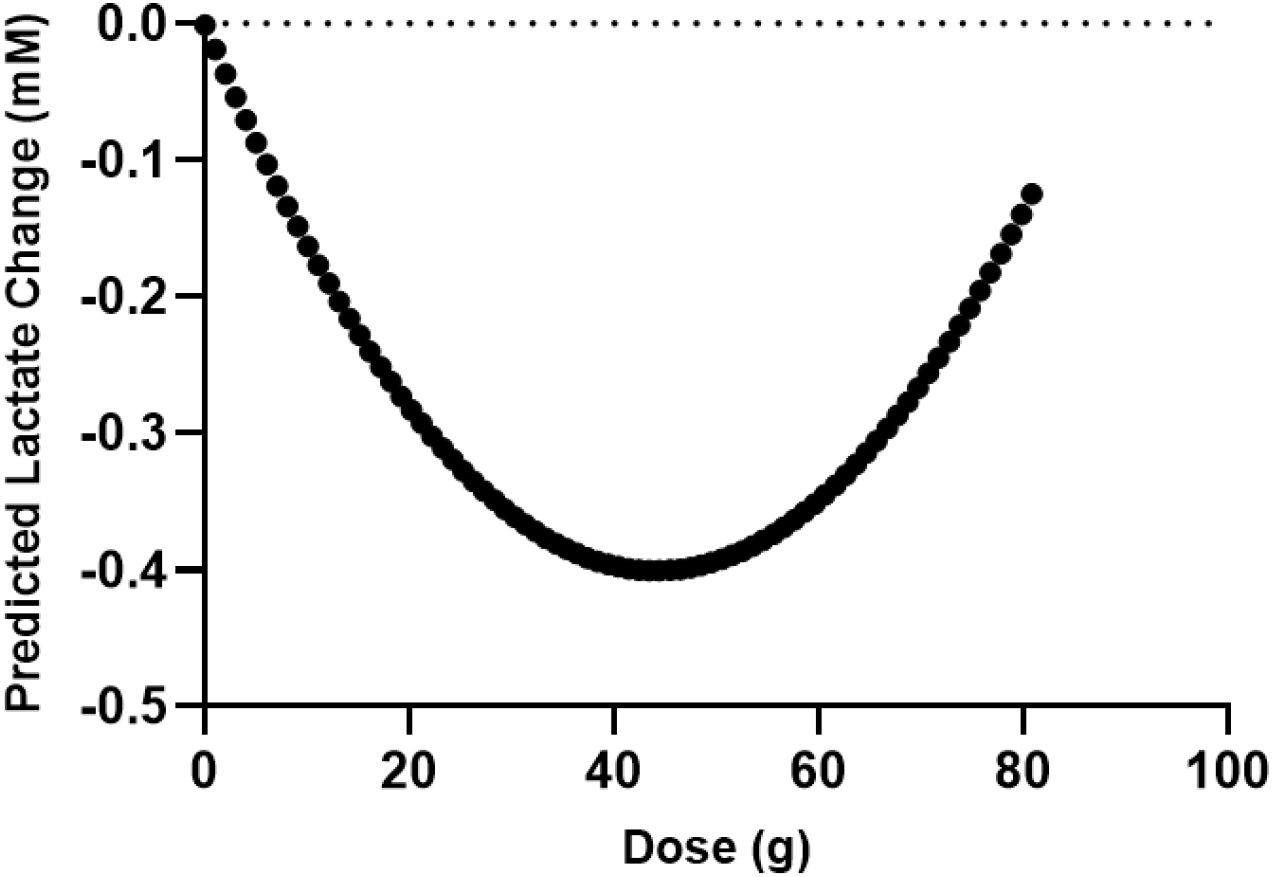
The quadratic regression model determined 44 g GRS as the optimal dose for reducing lactate accumulation.

### Optimal Dose Modeling

To identify the dose of GRS that results in the lowest blood lactate concentration during submaximal exercise, we applied a quadratic modeling approach. This analysis utilized data from three administered doses (Placebo, 37.5 g, and 75 g of GRS). The primary outcome variable was the change in blood lactate concentration at the highest submaximal work rate, using the placebo dose (0 g GRS) as a reference.

We incorporated baseline lactate levels measured in the morning before dosing, as an additional variable to account for inter-individual differences in lactate levels. A quadratic regression model was fitted to the data and the optimal dose was determined by calculating the vertex of the quadratic curve. This calculation identifies the dose at which the dose-response curve achieves its minimum value, representing the most effective dose for reducing lactate accumulation. The optimal dose for minimizing lactate accumulation was calculated as 44 g of GRS. To statistically evaluate the dose-response relationship, we employed a linear mixed-effects model. This model incorporated dose and baseline lactate levels as fixed effects and subject-specific variability as random effects to account for individual differences. The coefficients for the quadratic terms had an R^2^-value of 0.24 and p=0.025 confirming a U-shaped dose-response relationship.

## DISCUSSION

In the present study, we show that acute intake of a beverage based on GRS lowers blood lactate levels during submaximal cycling and alters blood glucose levels at rest and submaximal work. Intriguingly, we observed a decrease in lactate levels with both the 37.5 g and 75 g GRS compared to PLA during cycling close to the participants’ lactate threshold.

This result is consistent with our previous findings from the one-week GRS supplementation and HIIT-training study. In that trial, the results could not be explained by differences in either VO_2_max or mitochondrial function or capacity (Flockhart et al., 2023), indicating another unknown mechanism for changes in blood lactate and -glucose metabolism.

Animal studies on isothiocyanate supplementation have investigated the mechanisms underlying alterations in lactate metabolism following isothiocyanate supplementation. Wang et al. (2020) reported that SFN pretreatment in mice increased the protein levels of MCT1 in the skeletal muscle during acute hypoxic exercise. Since MCT1 facilitates the transport of lactate across cell membranes, this may explain the observed reduction in blood lactate levels. Additionally, the authors reported a significantly upregulated LDH activity which promoted the conversion of lactate to pyruvate (Wang et al., 2020). Overall, changes in the activity of the Cori cycle, which facilitates lactate clearance, represent an intriguing area for further investigation.

Clinical studies on dichloroacetate (DCA), a compound that stimulates pyruvate dehydrogenase (PDH), have demonstrated significantly reduced lactate levels during submaximal exercise and recovery, although not at maximal intensities. These findings suggest that limited PDH activity may play a role in the accumulation of plasma lactate during submaximal exertion in humans (Carraro et al., 1989). In the present study, the exercise intensity was moderate (RPE 10-15, Supplement, Fig. 2), and the modulation of PDH activity through isothiocyanate supplementation cannot be excluded as a contributing effect to the observed reductions in lactate levels.

Changes in the respiratory exchange ratio (RER) reflect shifts in substrate preference. A lower RER indicates increased fatty acid oxidation, whereas a higher RER reflects greater carbohydrate oxidation. Healthy individuals typically exhibit high metabolic flexibility, meaning their muscles can sense and utilize the most readily available substrate. Lower blood glucose levels signal a metabolic shift toward increased fat oxidation. Conversely, protons produced from the elevated production of lactic acid are buffered by the bicarbonate system and expelled as non-metabolic carbon dioxide in exhaled air, increasing the RER. In this study, we observed higher blood glucose levels at rest and during lower work rates, which diminished at the highest work rate. Conversely, blood lactate levels were significantly lower at the highest work rate but not at rest or the lowest work rate. Consistent with these findings, we observed a small but significant increase in RER at the lowest work rate following the 75 g dose, coinciding with the highest blood glucose levels. At the highest work rate, there was a significant reduction in RER after the 37.5 g dose, which coincided with the lowest lactate concentrations. In our previous study, we found a tendency towards a lower RER after one week of supplementation (Flockhart et al., 2023). Moreover, studies have demonstrated that mitochondrial oxidation of long-chain and short-chain fatty acids is depressed in the absence of Nrf2 and accelerated when Nrf2 is constitutively active (Ludtmann et al., 2014). Further studies on humans are required to confirm if SFN and/or activation of Nrf2 induces changes in substrate preference.

A limitation of this study is the difference in carbohydrate content between the beverages. The placebo was isocaloric to the 37.5 g dose, but the 75 g dose contained slightly more carbohydrates. However, the exercise tests were conducted three hours after consumption, in the postprandial state. Furthermore, we did not observe any differences in glucose concentrations between the 37.5 g and 75 g GRS doses, indicating that the difference in carbohydrate content was of minor importance. Another limitation is that we did not assess whether the lower lactate levels translated into improved performance. Theoretically, if the reduction in lactate results from the activation of pyruvate dehydrogenase, it could enhance performance by channeling more pyruvate into the mitochondria for efficient oxidation, reducing reliance on glycolysis, and thereby sparing muscle glycogen. Alternatively, if the lower lactate levels are due to increased activity of the hepatic Cori cycle, lactate could be more rapidly converted to glucose, possibly supporting glycogen resynthesis or maintaining blood glucose levels during exercise. Both mechanisms could potentially contribute to improved performance.

Isothiocyanates (ITCs) are electrophilic and have been shown to deplete endogenous glutathione (He et al., 2021) as well as induce single-strand DNA breaks (Sestili et al., 2010). This mild stress is thought to enhance endogenous antioxidant systems over the long term, partly via the activation of Nrf2. Notably, we observed a reduction in myeloperoxidase levels approximately three hours after GRS intake, suggesting a decrease in oxidative stress. This finding indicates that the adaptive compensatory system may act rapidly, likely within just a few hours of GRS consumption.

Blood glucose was significantly higher after both doses of GRS compared to PLA during rest and work rate 1. The increase in glucose levels could result from a lower oxidation of blood glucose or enhanced gluconeogenesis. However, during cycling at the lactate threshold, no significant differences in blood glucose were observed.

In conclusion, acute intake of small doses of GRS followed by submaximal ergometer cycling results in changes in lactate and glucose metabolism that could be beneficial for exercise performance.

## ACKNOWLEDGEMENT

Swedish Metabolomics Centre, Umeå, Sweden is acknowledged for metabolic profiling by LC-MS.

## SUPPLEMENTARY MATERIAL

**Supplementary material, Figure 1.**
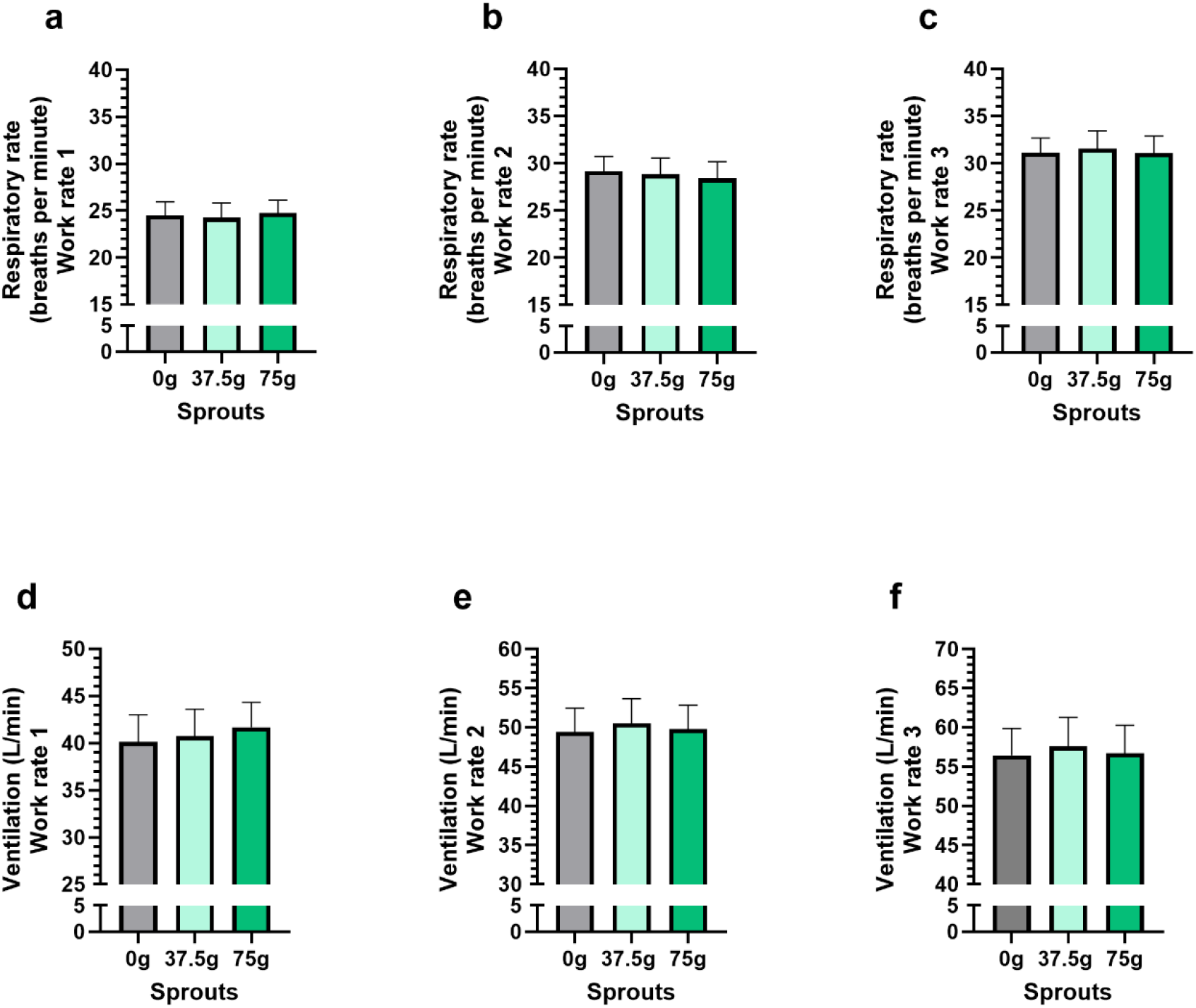
Respiratory rate (a-c) and ventilation (d-f) during cycling at 1-3 work rates on an ergometer bike, 3h after intake of a beverage containing 0 g, 37.5 g, or 75 g GRS. Data are presented as mean ± SEM, n = 15. p < 0.05 was considered significant.

**Supplementary material, Figure 2.**
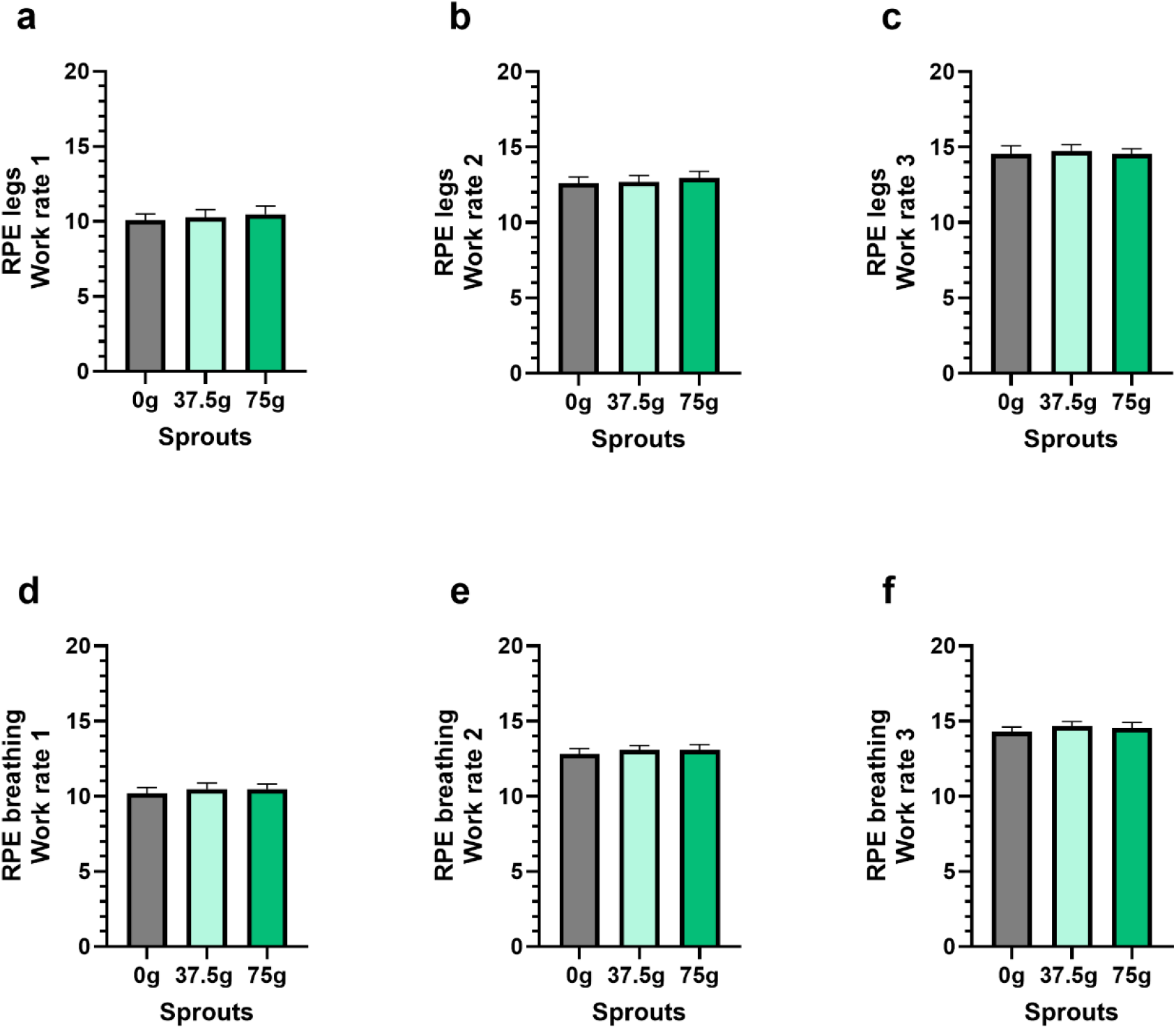
Rating of perceived exertion (RPE) in legs (a-c) and breathing (d-f) during cycling at 1-3 work rates on an ergometer bike, 3h after intake of a beverage containing 0 g, 37.5 g, or 75 g GRS. Data are presented as mean ± SEM, n = 15. p < 0.05 was considered significant.

## QUANTIFICATION OF ISOTHIOCYANATES

### Standards and Calibration Curve

Glucosinolate and isothiocyanate standards (L-sulforaphane, glucoraphin, glucoiberin and iberin) were obtained from Sigma-Aldrich (St. Louis, MO, USA). Stock solutions of each compound were prepared at a concentration of 1000 ng/μL and stored at -80ºC. An 11-point calibration curve (0.05-100 ng/µL) was prepared by serial dilutions and spiked with D6-salicylic acid as internal standards at a final concentration of 0.625 ng/μL.

### Sample Preparation

Sample preparation of 100µl was performed according to A et al 2005, with some modification. In detail, 900 µL of extraction buffer (90/10 v/v methanol: water) including internal standards (13C9-phenylalanine, D4-cholic acid, 13C9-caffeic Acid and D6-salicylic acid) was added to each sample together with 1 tungsten bead. The sample was shaken at 30 Hz for 3 minutes in a mixer mill. The sample was centrifuged at +4 °C, 14 000 rpm (18 620g), for 10 minutes. 150 or 200 µl supernatant was transferred to micro vials for LC-MS analysis.

A small aliquot of the remaining supernatants was pooled and used to create quality control (QC) samples. MSMS analysis was run on the QC samples for identification purposes. The samples were analyzed according to a randomized run order.

### LC-MS analysis

Before LC-MS analysis the sample was re-suspended in 10 + 10 µL methanol and water. All samples were first analyzed in positive mode, then the instrument was switched to negative mode and a second injection of each sample was performed.

The chromatographic separation was performed on an Agilent 1290 Infinity UHPLC-system (Agilent Technologies, Waldbronn, Germany). 2 μL of each sample were injected onto an Acquity UPLC HSS T3, 2.1 × 50 mm, 1.8 μm C18 column in combination with a 2.1 mm x 5 mm, 1.8 μm VanGuard precolumn (Waters Corporation, Milford, MA, USA) held at 40 °C. The gradient elution buffers were A (H2O, 0.1 % formic acid) and B (75/25 acetonitrile:2-propanol, 0.1 % formic acid), and the flow-rate was 0.5 mL min-1. The compounds were eluted with a linear gradient consisting of 0.1 - 10 % B over 2 minutes, B was increased to 99 % over 5 minutes and held at 99 % for 2 minutes; B was decreased to 0.1 % for 0.3 minutes and the flow-rate was increased to 0.8 mL min-1 for 0.5 minutes; these conditions were held for 0.9 minutes, after which the flow-rate was reduced to 0.5 mL min-1 for 0.1 minutes before the next injection.

The compounds were detected with an Agilent 6546 Q-TOF mass spectrometer equipped with a jet stream electrospray ion source operating in positive or negative ion mode. The settings were kept identical between the modes, with exception of the capillary voltage. A reference interface was connected for accurate mass measurements; the reference ions purine (4 μM) and HP-0921 (Hexakis(1H, 1H, 3H-tetrafluoropropoxy)phosphazine) (1 μM) were infused directly into the MS at a flow rate of 0.05 mL min-1 for internal calibration, and the monitored ions were purine m/z 121.05 and m/z 119.03632; HP-0921 m/z 922.0098 and m/z 966.000725 for positive and negative mode respectively. The gas temperature was set to 150°C, the drying gas flow to 8 L min-1 and the nebulizer pressure 35 psig. The sheath gas temp was set to 350°C and the sheath gas flow 11 L min-1. The capillary voltage was set to 4000 V in positive ion mode, and to 4000 V in negative ion mode. The nozzle voltage was 300 V. The fragmentor voltage was 120 V, the skimmer 65 V and the OCT 1 RF Vpp 750 V. The collision energy was set to 0 V. The m/z range was 70 - 1700, and data was collected in centroid mode with an acquisition rate of 4 scans s-1 (1977 transients/spectrum).

### Data analysis-evaluation/statistical methods

All data processing was performed using the Agilent Masshunter Profinder version B.10.0.2 (Agilent Technologies Inc., Santa Clara, CA, USA) using the Batch Targeted feature extraction in Masshunter Profinder and the glucosinolates as target library. The identification of the metabolites was based on MS, MSMS and retention time information. The concentration of each glucosinolate was calculated based on the calibration curves.

## REFERENCES

A, J., Trygg, J., Gullberg, J., Johansson, A. I., Jonsson, P., Antti, H., Marklund, S. L., & Moritz, T. (2005). Extraction and GC/MS analysis of the human blood plasma metabolome. Anal Chem, 77(24), 8086–8094. 10.1021/ac051211v

Billat, L. V. (1996). Use of blood lactate measurements for prediction of exercise performance and for control of training. Recommendations for long-distance running. Sports Med, 22(3), 157–175. 10.2165/00007256-199622030-00003

Calvert, L. D., Shelley, R., Singh, S. J., Greenhaff, P. L., Bankart, J., Morgan, M. D., & Steiner, M. C. (2008). Dichloroacetate enhances performance and reduces blood lactate during maximal cycle exercise in chronic obstructive pulmonary disease. Am J Respir Crit Care Med, 177(10), 1090–1094. 10.1164/rccm.200707-1032OC

Carraro, F., Klein, S., Rosenblatt, J. I., & Wolfe, R. R. (1989). Effect of dichloroacetate on lactate concentration in exercising humans. J Appl Physiol (1985), 66(2), 591–597. 10.1152/jappl.1989.66.2.591

Casado, A., Foster, C., Bakken, M., & Tjelta, L. I. (2023). Does Lactate-Guided Threshold Interval Training within a High-Volume Low-Intensity Approach Represent the “Next Step” in the Evolution of Distance Running Training? Int J Environ Res Public Health, 20(5). 10.3390/ijerph20053782

Fahey, J. W., Zhang, Y., & Talalay, P. (1997). Broccoli sprouts: an exceptionally rich source of inducers of enzymes that protect against chemical carcinogens. Proc Natl Acad Sci U S A, 94(19), 10367–10372. 10.1073/pnas.94.19.10367

Flockhart, M., Nilsson, L. C., Tillqvist, E. N., Vinge, F., Millbert, F., Lännerström, J., Nilsson, P. H., Samyn, D., Apró, W., Sundqvist, M. L., & Larsen, F. J. (2023). Glucosinolate-rich broccoli sprouts protect against oxidative stress and improve adaptations to intense exercise training. Redox Biol, 67, 102873. 10.1016/j.redox.2023.102873

He, C., Buongiorno, L. P., Wang, W., Tang, J. C. Y., Miceli, N., Taviano, M. F., Shan, Y., & Bao, Y. (2021). The Inhibitory Effect of Sulforaphane on Bladder Cancer Cell Depends on GSH Depletion-Induced by Nrf2 Translocation. Molecules, 26(16). 10.3390/molecules26164919

Hurley, B. F., Hagberg, J. M., Allen, W. K., Seals, D. R., Young, J. C., Cuddihee, R. W., & Holloszy, J. O. (1984). Effect of training on blood lactate levels during submaximal exercise. J Appl Physiol Respir Environ Exerc Physiol, 56(5), 1260–1264. 10.1152/jappl.1984.56.5.1260

Ludtmann, M. H., Angelova, P. R., Zhang, Y., Abramov, A. Y., & Dinkova-Kostova, A. T. (2014). Nrf2 affects the efficiency of mitochondrial fatty acid oxidation. Biochem J, 457(3), 415–424. 10.1042/bj20130863

Mawlong, I., Sujith Kumar, M. S., Gurung, B., Singh, K. H., & Singh, D. (2017). A simple spectrophotometric method for estimating total glucosinolates in mustard de-oiled cake. International Journal of Food Properties, 20(12), 3274–3281. 10.1080/10942912.2017.1286353

Oh, S., Komine, S., Warabi, E., Akiyama, K., Ishii, A., Ishige, K., Mizokami, Y., Kuga, K., Horie, M., Miwa, Y., Iwawaki, T., Yamamoto, M., & Shoda, J. (2017). Nuclear factor (erythroid derived 2)-like 2 activation increases exercise endurance capacity via redox modulation in skeletal muscles. Scientific Reports, 7(1), 12902. 10.1038/s41598-017-12926-y

Sestili, P., Paolillo, M., Lenzi, M., Colombo, E., Vallorani, L., Casadei, L., Martinelli, C., & Fimognari, C. (2010). Sulforaphane induces DNA single strand breaks in cultured human cells. Mutat Res, 689(1-2), 65–73. 10.1016/j.mrfmmm.2010.05.003

Sjödin, B., & Svedenhag, J. (1985). Applied physiology of marathon running. Sports Med, 2(2), 83–99. 10.2165/00007256-198502020-00002

Timmons, J. A., Gustafsson, T., Sundberg, C. J., Jansson, E., Hultman, E., Kaijser, L., Chwalbinska-Moneta, J., Constantin-Teodosiu, D., Macdonald, I. A., & Greenhaff, P. L. (1998). Substrate availability limits human skeletal muscle oxidative ATP regeneration at the onset of ischemic exercise. J Clin Invest, 101(1), 79–85. 10.1172/jci1146

Vargas-Mendoza, N., Morales-González, Á., Madrigal-Santillán, E. O., Madrigal-Bujaidar, E., Álvarez-González, I., García-Melo, L. F., Anguiano-Robledo, L., Fregoso-Aguilar, T., & Morales-Gonzalez, J. A. (2019). Antioxidant and Adaptative Response Mediated by Nrf2 during Physical Exercise. Antioxidants (Basel), 8(6). 10.3390/antiox8060196

Wang, L., Zhu, R., Wang, J., Yu, S., Wang, J., & Zhang, Y. (2020). Nrf2 Activation Enhances Muscular MCT1 Expression and Hypoxic Exercise Capacity. Medicine and science in sports and exercise, 52(8), 1719–1728. 10.1249/mss.0000000000002312

Wang, Y., Xiang, Y., Wang, R., Li, X., Wang, J., Yu, S., & Zhang, Y. (2022). Sulforaphane enhances Nrf2-mediated antioxidant responses of skeletal muscle induced by exhaustive exercise in HIIT mice. Food Science and Human Wellness, 11(5), 1355–1361. 10.1016/j.fshw.2022.04.035

